# Combinatorial Genetic Control of Rpd3S through histone H3K4 and H3K36 Methylation in Budding Yeast

**DOI:** 10.1101/376046

**Authors:** Kwan Yin Lee, Mathieu Ranger, Marc D. Meneghini

## Abstract

Much of euchromatin regulation occurs through reversible methylation of histone H3 lysine-4 and lysine-36 (H3K4me and H3K36me). Using the budding yeast *Saccharomyces cerevisiae*, we previously found that levels of H3K4me modulated temperature sensitive alleles of the transcriptional elongation complex Spt6-Spn1 through an unknown H3K4me effector pathway. Here we identify the Rpd3S histone deacetylase complex as the H3K4me effector underlying these Spt6-Spn1 genetic interactions. Exploiting these Spt6-Spn1 genetic interactions, we show that H3K4me and H3K36me collaboratively impact Rpd3S function in an opposing manner. H3K36me is deposited by the histone methyltransferase Set2 and is known to promote Rpd3S function at RNA PolII transcribed open reading frames. Using genetic epistasis experiments, we find that mutations perturbing the Set2-H3K36me-Rpd3S pathway suppress the growth defects caused by temperature sensitive alleles of *SPT6* and *SPN1*, illuminating that this pathway antagonizes Spt6-Spn1. Using these sensitive genetic assays, we also identify a role for H3K4me in antagonizing Rpd3S that functions through the Rpd3S subunit Rco1, which is known to bind H3 N-terminal tails in a manner that is prevented by H3K4me. Further genetic experiments reveal that the H3K4 and H3K36 demethylases *JHD2* and *RPH1* mediate this combinatorial control of Rpd3S. Finally, our studies also show that the Rpd3L complex, which acts at promoter-proximal regions of PolII transcribed genes, counters Rpd3S for genetic modulation of Spt6-Spn1, and that these two Rpd3 complexes balance the activities of each other. Our findings present the first evidence that H3K4me and H3K36me act combinatorially to control Rpd3S.

## Introduction

Unstructured N-terminal tails of histone proteins emanate from the nucleosome core and are sites of a diverse array of post-translational modifications (Strahl and Allis 2000). Methylations of histone H3 on lysine-4 and lysine-36 (H3K4me and H3K36me) are among the most well studied histone post-translational modifications. Like all lysine methylations, H3K4 and H3K36 can be mono, di, or tri methylated (me1, me2, me3), and these forms have distinctive roles in regulation of proteins with domains that distinguish these states (Yun *et al*. 2011; Musselman *et al*. 2012; Rando 2012). Together with a plethora of other histone post-translational modifications, H3K4me and H3K36me contribute to a diverse chromatin landscape. A prediction of the histone code hypothesis states that chromatin effector complexes interpret this diverse landscape through their ability to distinguish multiple histone modifications combinatorially (Strahl and Allis 2000). Indeed, chromatin effector complexes often contain multiple subunits with protein domains known to distinguish histone modification states (Doyon and Cote 2004; Li *et al*. 2007; Wang *et al*. 2011). However, although this prediction has been satisfied with clear biochemical support (Li *et al*. 2007; McDaniel *et al*. 2016), there exists little genetic evidence specifically addressing this component of the histone code hypothesis.

Rpd3 is the catalytic subunit of the conserved histone deacetylase complexes Rpd3L and Rpd3S in the yeast *S. cerevisiae* (Rundlett *et al*. 1996; Bernstein *et al*. 2000; Kurdistani *et al*. 2002). Distinctive functions of Rpd3S and Rpd3L are enabled by their chromatin recruitment to discrete regions of RNA PolII transcribed genes, with Rpd3L being recruited to 5’ promoter-proximal regions (Carrozza *et al*. 2005a) while Rpd3S is principally found downstream within the gene bodies (Carrozza *et al*. 2005b; Keogh *et al*. 2005). Both Rpd3S and Rpd3L share the three core subunits Rpd3, Sin3, and Ume1 and their differing localizations on chromatin are likely facilitated through additional protein subunits that distinguish these complexes. Rpd3L contains numerous additional protein subunits (Carrozza *et al*. 2005a), and controls meiotic progression in diploid cells as well as the transcriptional program of haploid cells entering quiescence (Lardenois *et al*. 2015; McKnight *et al*. 2015). The Rco1 and Eaf3 proteins are specific for Rpd3S, which negatively regulates transcriptional elongation by RNA PolII in mitotically proliferating cells and prevents spurious intergenic transcription (CARROZZA *et al*. 2005b; Joshi and Struhl 2005; Keogh *et al*. 2005; Quan and Hartzog 2010). Eaf3 possesses a biochemically confirmed capacity for binding to H3K36me (LI *et al*. 2007; Xu *et al*. 2008; Ruan *et al*. 2015; Steunou *et al*. 2016) and accordingly mediates Rpd3S activity at RNA PolII transcribed open reading frames where H3K36me is found (Carrozza *et al*. 2005b; Keogh *et al*. 2005). The function of Rpd3S also requires the Rco1 subunit, which binds the H3 N-terminal tail through its tandem PHD domains (Lee *et al*. 2013; Ruan *et al*. 2015; McDaniel *et al*. 2016). Interestingly, methylation of H3K4 prevents Rco1 binding to the H3 N-terminus *in vitro* (McDaniel et al. 2016), though the *in vivo* roles of H3K4me for Rpd3S function remain unknown.

We previously showed that the Spt6-Spn1 histone chaperone complex was genetically governed in an H3K4me3-dependent manner through the opposing roles of the H3K4 methyltransferase and demethylase enzymes Set1 and Jhd2 (LEE *et al*. 2018). Mutations in *SET1* and other genes that cause reduced H3K4me3 enhance the growth defects caused by temperature sensitive alleles of *SPN1* and *SPT6*, while mutation of *JHD2*, which causes increased H3K4me3, suppress Spt6-Spn1. These and other results supported our conclusion that H3K4me3 levels impacted Spt6-Spn1, though the pathways underlying Spt6-Spn1 regulation by H3K4me remained opaque (Lee *et al*. 2018).

Here we identify Rpd3S as the H3K4me effector pathway impacting Spt6-Spn1 and provide genetic evidence for the biochemically predicted, though untested, model that H3K4me collaborates with H3K36me to combinatorially control the function of Rpd3S through its Eaf3 and Rco1 subunits. We find that mutating components throughout the Set2-H3K36me-Rpd3S pathway suppress Spt6-Spn1 mutations, suggesting that activation of Rpd3S through H3K36me opposes Spt6-Spn1. Using sensitive epistasis experiments, we show that in opposition to this known H3K36me Rpd3S activating role, H3K4me negatively impacts Rpd3S. Further genetic experiments suggest that H3K4me opposes Rpd3S by inhibiting Rco1 binding to H3 N-terminal tails. Our genetic findings are in good agreement with the biochemically characterized specificities of the chromatin binding domains within Rco1 and Eaf3, and suggest that these binding specificities fine-tune the action of Rpd3S on chromatin.

## Materials and Methods

### Yeast Strains

Standard yeast genetic methods were used for construction of all strains and deletion mutants were obtained from the gene deletion collection (Giaever *et al*. 2002). The GAL inducible *JHD2* alleles were constructed using PCR based integration as described (Longtine *et al*. 1998). The *JHD2(H427A)* mutation was constructed using the delitto perfetto method (Storici and Resnick 2003). Yeast strains used in this study are listed in Table S1. All strains were constructed through genetic crosses followed by dissections in the BY4742 background.

### Serial dilution assays

Yeast strains were inoculated into several mL of YPD (1% yeast extract, 2% peptone, and 2% glucose) and grown overnight at room temperature (23°C). Each strain was diluted to an 0D600 = 0.4, then 10-fold serially diluted five times and spotted on synthetic complete (SC) media (YNB media (Multicell Wisent) containing 5 g/L of ammonium sulfate and either 2% glucose or 2% galactose as described previously (Lee *et al*. 2018).

### Western blots

Cells were grown to mid-logarithmic phase in synthetic complete (SC) medium + 2% raffinose and transferred to SC + 2% galactose medium. Aliquots of these cultures were taken 1 hour after transfer to galactose-containing media. Proteins were then extracted and processed for western blotting as described previously (Xu *et al*. 2012). Equal amounts of protein were electrophoresed on SDS-PAGE gels and transferred onto PVDF membranes. Immunoblot analysis was performed as described (Soloveychik *et al*. 2016). Visualization of the proteins was performed by exposing the membrane to light sensitive film. The PVDF membrane was stripped and re-probed with different antibodies. Stripping the PVDF membrane was accomplished by incubating the membrane for 30 minutes at 50° C with a mixture consisting of 100 mM 2-β-mercaptoethanol, 2% SDS and 62.5 mM Tris-HCl pH 6.7.

### Antibodies

The following antibodies were used in this study and purchased from Abcam: Anti-Histone H3 antibody (ab1791), Anti-Histone H3K4me3 antibody (ab8580), and Anti-Histone H3K4me1 antibody (ab8895). The following antibody was used in this study and purchased from Millipore: Anti-Histone H3K4me2 (07-030).

## Results

### Opposing roles of H3K4me and H3K36me in control of Spt6-Spn1

We previously found that deletion of *JHD2* (*jhd2*Δ) suppressed TS alleles of *SPT6* and *SPN1* and this was attributable to increased H3K4me levels caused by *jhd2*Δ (Lee et al. 2018). Because *set2*Δ was previously found to suppress TS alleles in the functionally related FACT complex (Biswas *et al*. 2006), we asked if *set2*Δ similarly suppressed Spt6-Spn1 TS mutations. We found that *set2*Δ strongly suppressed the temperature sensitive growth defects caused by the *spt6-14* and *spn1-K192N* mutations (Figure 1A and 1B). The magnitude of suppression caused by *set2*Δ was greater than that caused by *jhd2*Δ and no additive effects on suppression were observed (Figure 1A and 1B). All strains described here were engineered using genetic crosses and tetrad dissection. For all results shown, we isolated at least 2 independently constructed strain replicates through tetrad dissection. Though replicates and WT control strains are not always shown in the interest of space, all results we report here are upheld in these replicates. strain fitness was assessed using spot assays to compare growth rates at varied temperatures. Though also not shown in the interest of space, we always observed complete loss of growth of our TS strains at 38.5°C, confirming that the TS alleles were intact and no bypass suppression occurred.

**Figure 1.**
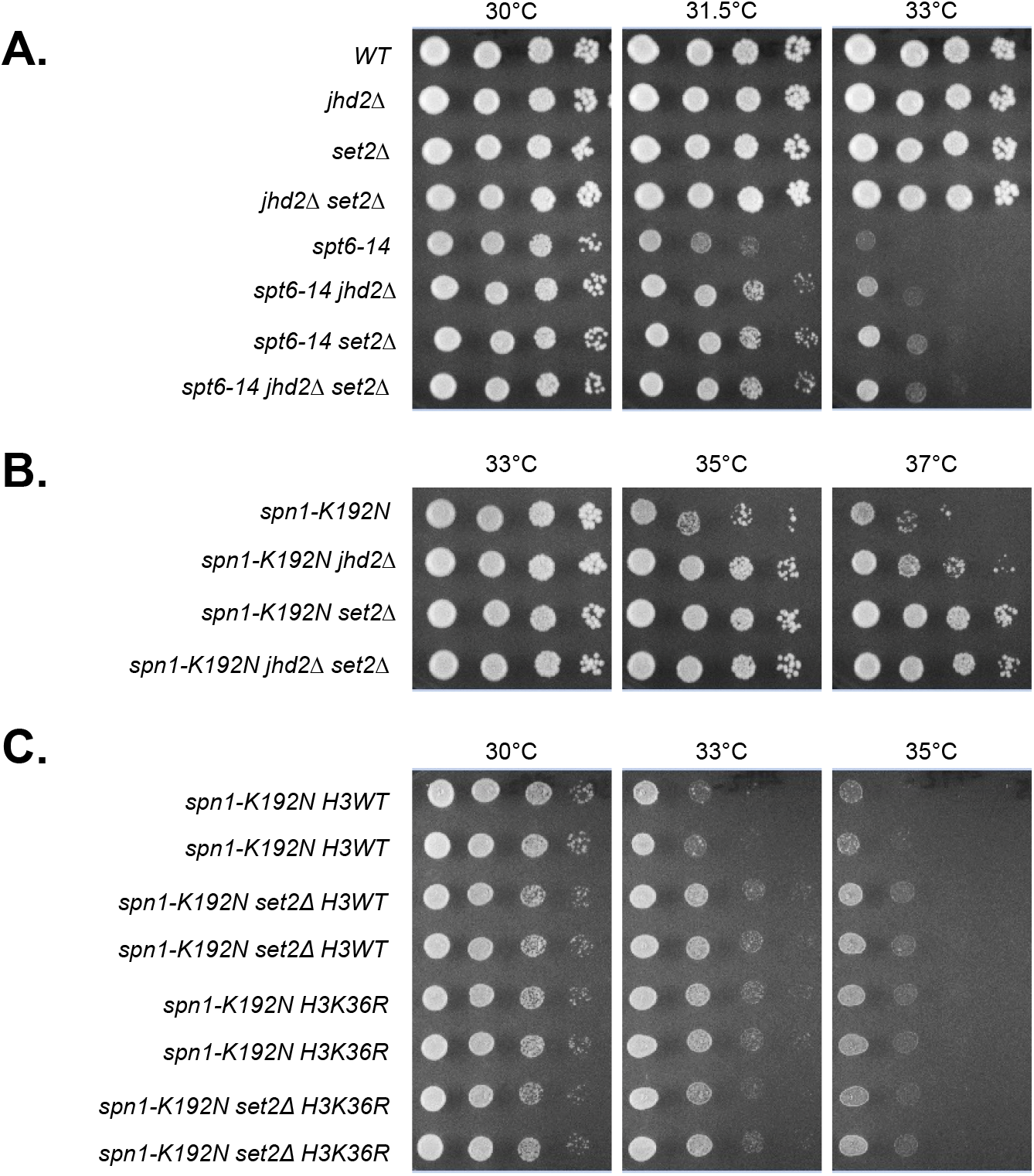
Set2 antagonizes Spt6-Spn1 through H3K36me. Yeast strains with the indicated genotypes were serially diluted ten-fold, spotted onto agar plates containing synthetic complete media, and grown at indicated temperatures. Genetic interactions of *jhd2*Δ and *set2*Δ with the temperature sensitive alleles (A) *spt6-14* and (B) *spn1-K192N* are shown. (C) Strains were constructed through genetic crosses with H3K36R substitution mutants from the Dharmacon Non Essential Histone H3 & H4 Mutant Collection. Genetic interactions of *set2*Δ and H3K36R with *spn1-K192N* are shown.

As the only known substrate of Set2 is H3K36, we hypothesized that the loss of H3K36me caused by *set2*Δ accounted for the suppression of Spt6-Spn1 TS alleles. To test this, we engineered strains combining *spn1-K192N* with synthetic histone H3 alleles encoding histone H3 lysine 36 substituted with arginine (H3K36R) (Dai *et al*. 2008). The H3K36R substitution suppressed *spn1-K192N* temperature sensitivity compared with an isogenic strain expressing a wild type synthetic histone allele (Figure 1C). The magnitude of suppression by H3K36R substitution was equivalent to that of *set2*Δ and their combined influence was not additive (Figure 1C). This result supports the conclusion that suppression of *spn1-K192N* by *set2*Δ occurred through loss of H3K36me, showing that H3K36me somehow opposed Spn1 function.

To genetically deconvolute the impact of H3K4me and H3K36me on Spt6-Spn1, we engineered strains expressing the endogenous *JHD2* gene under control of the *GAL1-10* promoter *(P_GAL_-JHD2)*, which leads to *JHD2* overexpression in galactose medium. We found that growth of strains harboring *P_GAL_-JHD2* in galactose containing media caused dramatic depletions of H3K4me3 and H3K4me2 (Figure 2A). As expected based on our previous findings, on media containing glucose, which leads to strong transcriptional repression of *GAL1-10, P_GAL_-JHD2* phenocopied *jhd2*Δ and caused suppression of both *spt6-14* and *spn1-K192N* (Figure 2B and data not shown). On media containing galactose however, we found that overexpression of *JHD2* negatively affected the growth of *spt6-14* and *spn1-K192N* (Figure 2C and 2D), consistent with our previous finding that mutations reducing H3K4me3 enhanced these Spt6-Spn1 alleles (Lee *et al*. 2018). Indeed, the enhancement of Spt6-Spn1 TS mutations caused by *P_GAL_-JHD2* in the presence of galactose was reverted by mutation of Jhd2 histidine-427 to alanine (H427A), which renders Jhd2 catalytically inactive and unable to demethylate H3K4 (Ingvarsdottir *et al*. 2007; LIANG *et al*. 2007) (Figure 2A and S1). Using the *P_GAL_-JHD2* allele, we found that suppression of both *spt6-14* and *spn1-K192N*by *set2*Δ was epistatic to the growth defect caused by *JHD2* overexpression (Figure 2C and Figure 2D). To confirm that this epistatic relationship related specifically to H3K4 demethylation by Jhd2, we determined that *set2*Δ similarly suppressed the phenotypic enhancement of *spn1-K192N* caused by *setl*Δ (Figure 2E).

**Figure 2.**
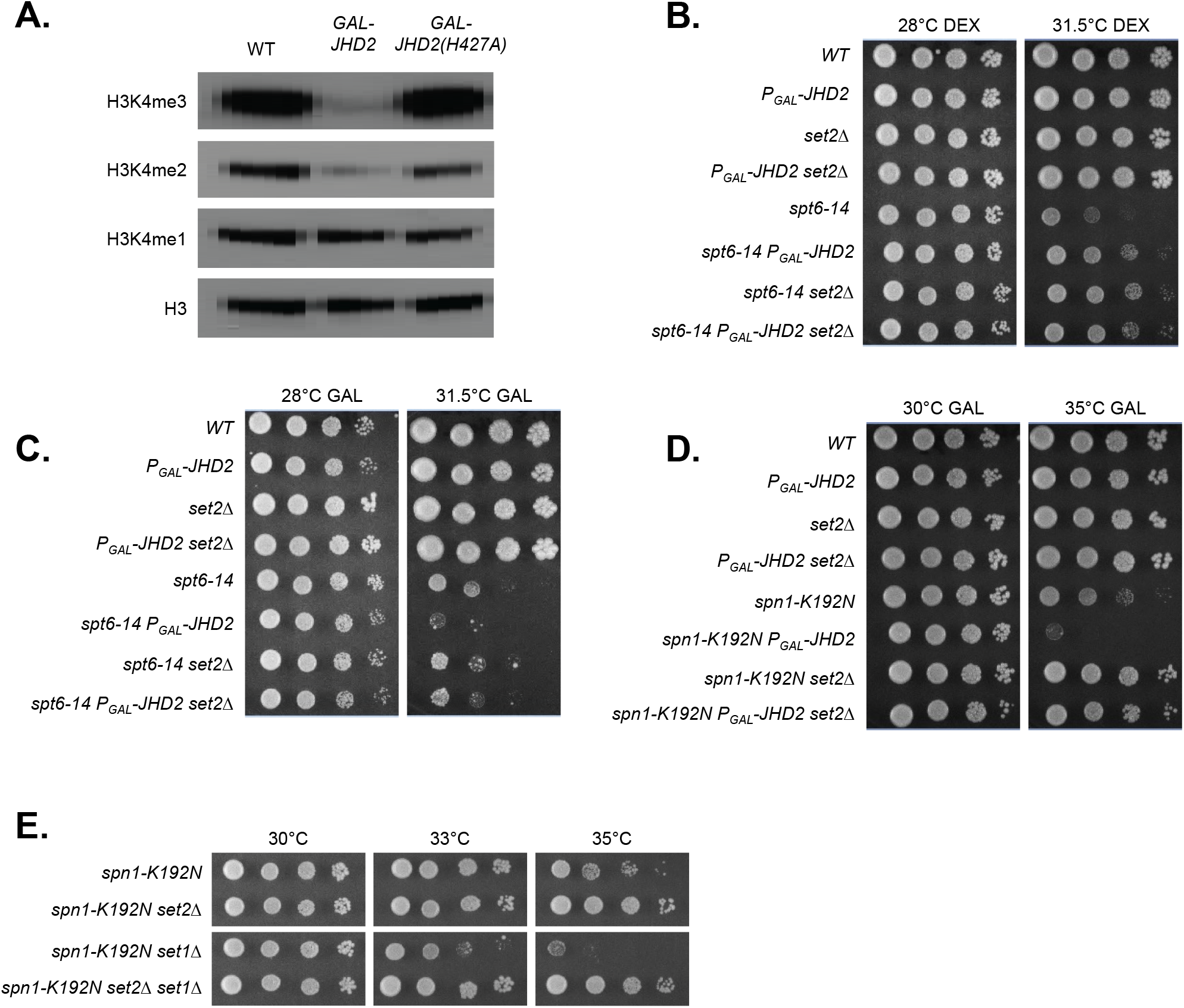
Suppression of Spt6-Spn1 mutants by *set2*Δ is epistatic to their enhancement caused by *JHD2* overexpression. *P_GAL_-JHD2* is a construct that replaces the endogenous *JHD2* locus. Cells grown in galactose overexpress *JHD2* and cells grown without galactose do not express *JHD2*. (A) Western blot detection of H3K4me3, H3K4me2, H3K4me1, and pan-H3 are shown for extracts prepared from wild-type, *P_GAL_-JHD2*, and *P_GAL_-JHD2(H427A)* cells grown in synthetic complete media containing 2% galactose. Pan-H3 serves as a loading control. (B, C, and D) Plate spot assays (as described in Figure 1) were used to compare the growth of the indicated strains on synthetic media with 2% dextrose (DEX, B) or 2% galactose (GAL, C and D). (E) Genetic interactions of *set1*Δ and *set2*Δ with *spn1-K192N* are shown.

Collectively, these results demonstrate that Spt6-Spn1 activity was opposed both by methylation of H3K36 and by hypo-methylation of H3K4 (denoted hereafter as “H3K4me0”). Moreover, the epistatic relationships we show suggest that the methylation state of H3K36 had a more crucial role than that of H3K4 in Spt6-Spn1 regulation. The remainder of the work described here exploits the Spt6-Spn1 TS mutations as a tool enabling genetic interrogation of this prospective H3K4me0/H3K36me regulatory pathway. Because *spn1-K192N* generally provided more robust genetic interactions compared with *spt6-14*, most of our studies used the *spn1-K192N* mutation and we show these here, though we always observed qualitatively equivalent interactions using *spt6-14*.

### H3K4 and H3K36 methylation states collaboratively modulated Rpd3S

A parsimonious model explaining our results posits an effector complex that opposes Spt6-Spn1 which itself is combinatorially controlled by H3K4/36me states. The Rpd3S histone deacetylase complex possesses two requisite characteristics that satisfy this hypothesis: 1, Rpd3S is positively regulated through an interaction of its Eaf3 subunit with H3K36me (Li *et al*. 2007; Xu *et al*. 2008; Ruan *et al*. 2015); and 2, The Rco1 subunit of Rpd3S binds to H3 N-terminal tails, and methylation of H3K4 opposes Rco1 binding (Lee *et al*. 2007; Ruan *et al*. 2015; McDaniel *et al*. 2016). We determined that the temperature sensitive growth defects caused by *spn1-K192N* were suppressed by *rpd3*Δ and that suppression by *rpd3*Δ and *jhd2*Δ were not additive, similarly to what we found with *jhd2*Δ and *set2*Δ (Figure 3A). Supporting the model that a loss of Rpd3S function specifically accounted for this result, we found that *rco1*Δ and *eaf3*Δ equivalently suppressed *spn1-K192N* (Fig 3B and data not shown). Eaf3 is also found in the NuA4 histone acetyltransferase complex. To rule out a role for Eaf3 as a component of NuA4 in our findings, we interrogated genetic interactions of *eaf7Δ*, which causes a specific loss of Eaf3 from NuA4 while leaving Eaf3 incorporation into Rpd3S unperturbed (Rossetto *et al*. 2014). Deletion of *EAF7* caused no detectable genetic interactions with *spn1-K192N*, further upholding the conclusion that perturbation of Rpd3S specifically accounted for our findings (Figure S2).

**Figure 3.**
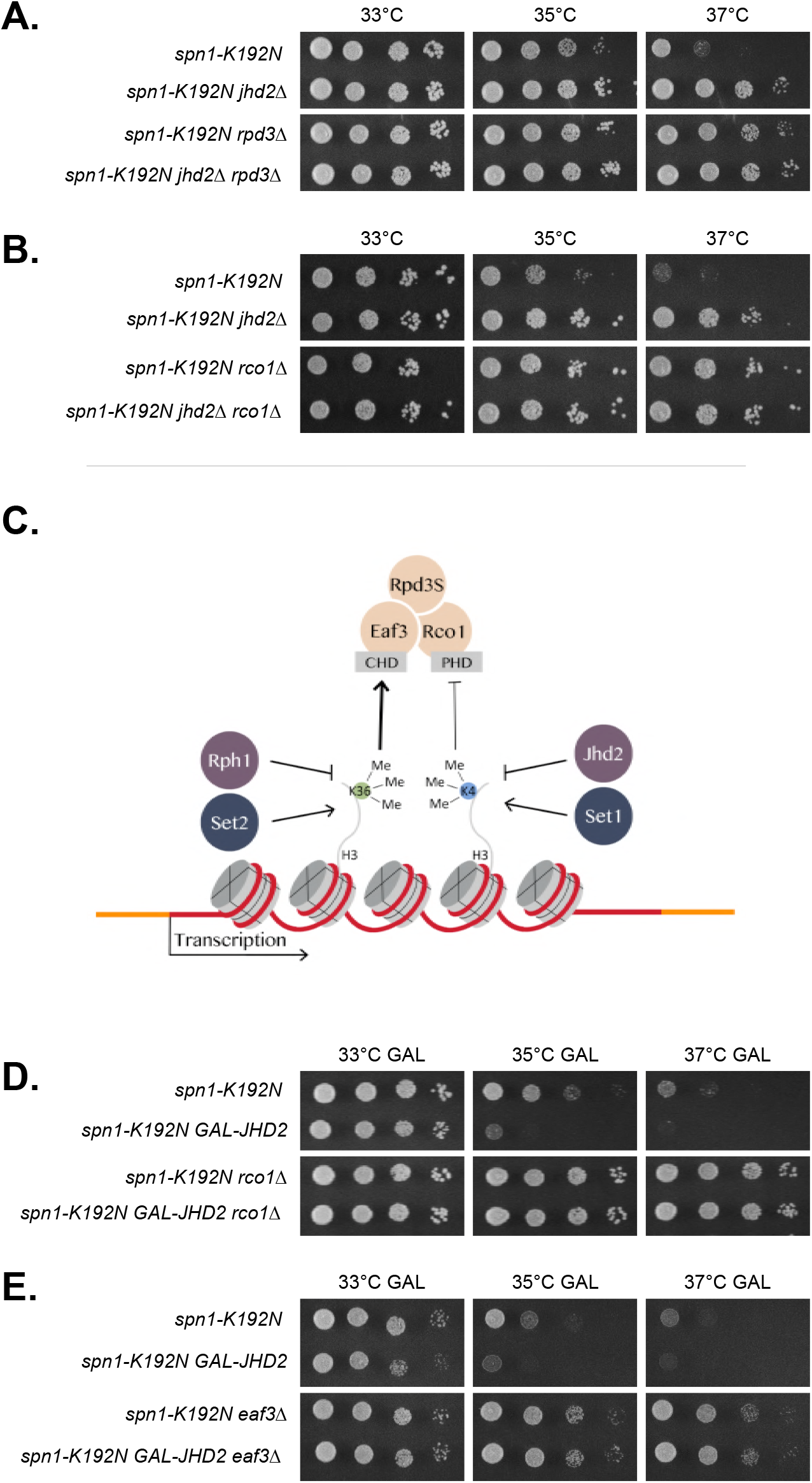
*SPN1* mutants were suppressed by loss of Rpd3S. Plate spot assays (as described in Figure 1) were used to compare the growth of the indicated strains on synthetic media with 2% dextrose (DEX) or 2% galactose (GAL). (A) Genetic interactions of *jhd2*Δ and *rpd3*Δ with *spn1-K192N*. (B) Genetic interactions of *jhd2*Δ and *rco1*Δ with *spn1-K192N. P_GAL_-JHD2* (as described in Figure 2) is used in the following experiments. (C) Genetic model of Rpd3S regulation by the H3K4me and H3K36me through the methyl-binding specificities of the Rpd3S subunits Rco1 and Eaf3. The heavier arrow indicates that H3K36me has a more crucial role in the regulation of Rpd3S than that of H3K4me0. (D) Genetic interactions of *rco1*Δ and *P_GAL_-JHD2* with *spn1-K192N* on GAL. (E) Genetic interactions of *eaf3*Δ and *P_GAL_-JHD2* with *spn1-K192N* on GAL.

The activation of Rpd3S by H3K36me is well established. We propose that H3K4me0 acts collaboratively with H3K36me, though in a comparatively minor fashion, to promote Rpd3S function (Figure 3C). To test this, we again employed *P_GAL_-JHD2* strains. Like *set2Δ*, suppression of *spn1-K192N*by both *eaf3*Δ and *rco1*Δ was epistatic to the enhanced growth defect caused by *JHD2* galactose overexpression (Figure 3D and 3E). As mutations in *RCO1* and *EAF3* disrupt Rpd3S while leaving the Rpd3L complex intact (Carrozza *et al*. 2005b), these findings are consistent with Rpd3S mediating the proposed H3K4me0/H3K36me effector pathway (Figure 3C).

In contrast to the well-characterized mechanistic role of H3K36me in activation of Rpd3S through Eaf3, the potential role of H3K4me in modulation of Rpd3S through Rco1 is unexplored. Rco1 binds to H3 N-terminal tails that are hypo-methylated on H3K4 through its tandem PHD domains, each of which is essential for interaction with the H3 N-terminus (McDaniel *et al*. 2016). To specifically evaluate the genetic consequences of loss of H3K4 binding by Rco1, we made use of an allele of Rco1 that lacks one of these PHD domains *(rco1-PHDΔ)* but encodes a stable protein that is assembled into Rpd3S *in vivo* (LI et al. 2007). We found that *rco1-PHD*Δ suppressed *spn1-K192N* equivalently to *jhd2Δ*, and that these phenotypes were not additive (Figure 4A). These results support our model that Jhd2 opposed Rpd3S function by increasing the methylation state of H3K4, thereby restricting the ability of Rco1 to activate Rpd3S through its H3 N-terminal tail binding activity (Figure 3C). A further prediction of this model is that *rco1*Δ should also suppress the enhanced *spn1-K192N*growth defects caused by the complete loss of H3K4me (Lee *et al*. 2018). We tested this using the H3K4R allele and found this was indeed the case: *rco1*Δ suppressed the growth defects of *spn1-K192NH3K4R* mutants (Figure 4B).

**Figure 4.**
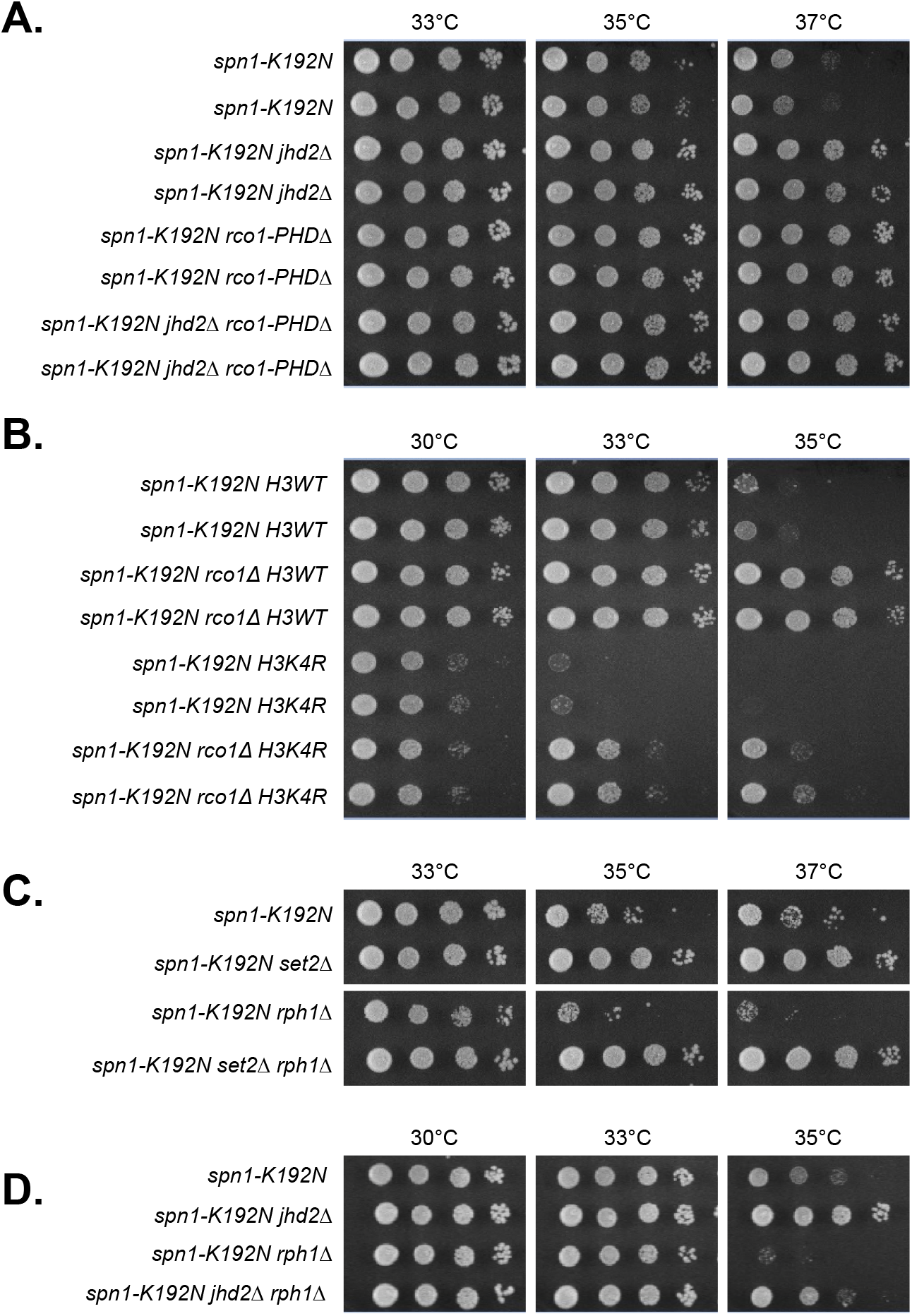
H3K4 and H3K36 methylation states collaboratively modulated Rpd3S. Plate spot assays (as described in Figure 1) were used to compare the growth of the indicated strains. *rco1-PHD*Δ is an allele of *RCO1* that lacks the N-terminal PHD domain but encodes a stable protein that is assembled into Rpd3S *in vivo*. (A) Genetic interactions of *jhd2*Δ and *rco1-PHD*Δ with *spn1-K192N*. Growth of two independent isolates of each genotype is shown. (B) Genetic interactions of *rco1*Δ and H3K4R with *spn1-K192N*. Growth of two independent isolates of each genotype is shown. (C) Genetic interactions of *rphl*Δ and *set2*Δ with *spn1-K192N*. (D) Genetic interactions of *jhd2*Δ and *rphl*Δ with *spn1-K192N*.

Like H3K4me, H3K36me is also subject to the opposing roles of methyltransferases and demethylases. Our model predicts that increased H3K36me levels caused by loss of H3K36 demethylation should show the opposite phenotype as *set2*Δ and enhance the temperature sensitive growth defect of *spn1-K192N*. Yeast possess two confirmed H3K36 demethylases belonging to the Jumonji superfamily, Jhd1 and Rph1. Rph1 demethylates both H3 K36 tri- and dimethyl substrates (Kim and Buratowski 2007; Klose *et al*. 2007) while Jhd1 targets di- and monomethyl H3K36 (Tsukada *et al*. 2006; Fang *et al*. 2007). We found no consequence for *jhdl*Δ in any of our experiments (data not shown). In contrast to *jhd1Δ*, we found that *rphl*Δ strongly exacerbated the *spn1-K192N* growth defects, and that this consequence was reverted by *set2Δ*, suggesting that Rph1-mediated H3K36 demethylation opposed Rpd3S function (Figure 3C and 4C). Indeed, *spn1-K192N*enhancement by *rphl*Δ was also reverted by *rco1*Δ (Figure S3). Taking into consideration the biochemical activities of Jhd1 and Rph1, our findings suggest that it is the tri-methylated state of H3K36 (H3K36me3) that specifically activates Rpd3S, consistent with biochemical characterization of Eaf3 binding specificity (Steunou *et al*. 2016). Finally, we found that *spn1-K192N* enhancement by *rph1*Δ was partially suppressed by *jhd2*Δ (Figure 4D). The partial suppression of *spn1-Kl92Nrphl*Δ by *jhd2*Δ is consistent with the more predominant role we propose for H3K36me in Rpd3S activation compared with H3K4me0 (Figure 3C).

### Rpd3L complex availability counterbalanced Rpd3S function

Surprisingly, we found that *rpd3*Δ suppressed *spn1-K192N*less robustly than *rcol*Δ, and that this less robust suppression was in fact epistatic to *rco1Δ*, suggesting that loss of other Rpd3 functions had detrimental effects on *spn1-K192N* (Figure 5A). The Rpd3L complex seemed like the best candidate for such a counterbalancing Rpd3 function. To assess the function of Rpd3L in our experiments, we utilized mutations in Rpd3L subunits that had varying effects on Rpd3L complex formation. Loss of the Pho23 subunit of Rpd3L is documented to have only minor consequences on complex integrity (CARROZZA *et al*. 2005a). *PHO23* deletion had no effect on *spn1-K192N* temperature sensitivity nor did it affect suppression of *spn1-K192N* by *jhd2*Δ (Figure 5B). In contrast to *PHO23*, we found that *depl*Δ, which was previously shown to completely disrupt Rpd3L complex formation (Carrozza *et al*. 2005a), enhanced the temperature sensitive growth defect of *spn1-K192N* (Figure 5C). Thus, Rpd3L appeared to somehow promote *SPN1* function in direct contrast to its Rpd3S counterpart.

**Figure 5.**
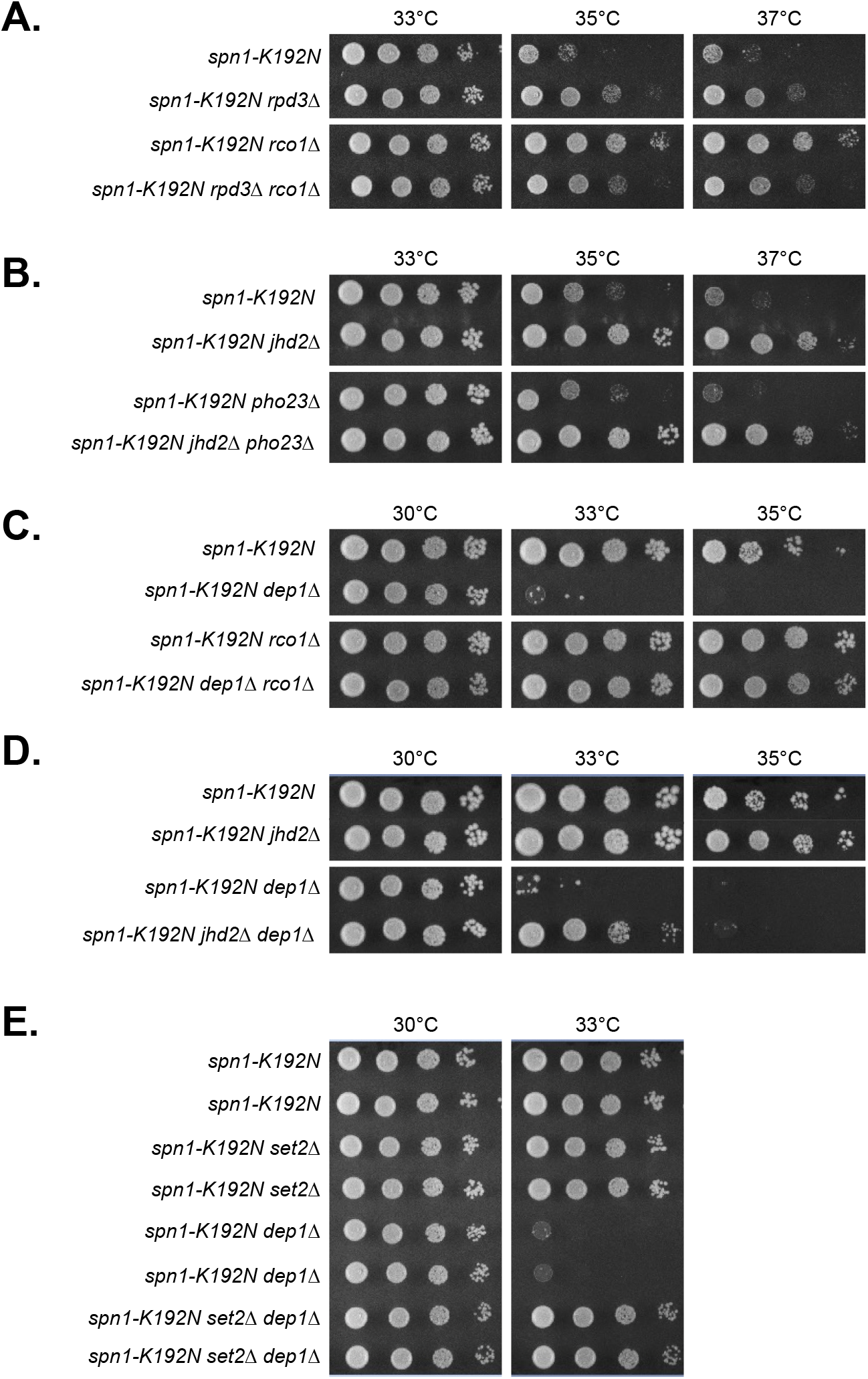
Rpd3L complex availability counterbalanced Rpd3S function. Plate spot assays (as described in Figure 1) were used to compare the growth of the indicated strains. (A) Genetic interactions of *rcol*Δ and *rpd3*Δ with *spn1-K192N*. (B) Genetic interactions of *jhd2*Δ and *pho23*Δ with *spn1-K192N*. (C) Genetic interactions of *depl*Δ and *rcolΔ*, with *spn1-K192N*. (D) Genetic interactions of *depl*Δ and *jhd2*Δ with *spn1-K192N*. (E) Genetic interactions of *depl*Δ and *set2*Δ with *spn1-K192N*. Growth of two independent isolates of each genotype is shown.

The disruption of Rpd3L was previously shown to greatly increase the amount of Rco1 protein in the cell (Biswas *et al*. 2008). We hypothesized that *depl*Δ enhanced the temperature sensitivity of *spn1-K192N* via increased Rco1 protein levels and resulting increased Rpd3S function. To test this, we introduced *rco1*Δ into the *spn1-K192Ndepl*Δ double mutant. Suppression of *spn1-K192N*by *rcol*Δ was epistatic to the enhancement caused by *depl*Δ suggesting that the *depl*Δ enhancement phenotype was due to increased Rpd3S function (Figure 5C). As our model posits that Jhd2 positively regulated Rpd3S through its demethylation of H3K4, *jhd2*Δ is predicted to therefore also alleviate the enhanced *spn1-K192N* growth defects caused by loss of Rpd3L. Consistent with this prediction, we found that *jhd2*Δ suppressed the growth defects of *spn1-K192N depl*Δ mutants, but not to the same extent as *rco1*Δ did (Figure 5D). This less robust suppression of *spn1-K192N*by *jhd2*Δ compared with by *rcol*Δ seems sensible, as Rpd3S is presumably still present in *jhd2*Δ cells while it is completely absent in *rcol*Δ cells (Carrozza *et al*. 2005a). As expected given the known role of Set2 in promoting Rpd3S chromatin recruitment, *set2Δ*, like *rco1Δ*, reverted the *spn1-Kl92Ndepl*Δ growth defects. (Figure 5E).

## Discussion

We previously found that increased H3K4me3 levels suppressed TS alleles of Spt6-Spn1 while decreased H3K4me3 levels enhanced them (Lee *et al*. 2018). Here we show that perturbation of the Set2-H3K36me-Rpd3S pathway similarly suppresses TS alleles of Spt6-Spn1, revealing that Rpd3S function somehow opposes Spt6-Spn1 and permitting a series of genetic epistasis experiments investigating the contributions of H3K4me and H3K36me to Rpd3S functionality. Indeed, we provide genetic evidence consistent with the conclusion that H3K4me also modulates Rpd3S: *rco1*Δ and *rco1-PHD*Δ suppressed *spn1-K192N* equivalently to and non-additively with *jhd2*Δ. Critically, *rco1*Δ also suppressed the exacerbated *spn1-K192N* growth defects caused by reduced H3K4me. Our subsequent genetic interrogations of both *RPH1* and of Rpd3L further support the model we present in Figure 3C. Functional insights into H3K4 and H3K36 demethylation in yeast remain relatively narrow compared with their corresponding methyltransferase enzymes, and our findings show that at least one role of Jhd2 and Rph1 is in fine-tuning H3K4me0/H3K36me3 activation of Rpd3S.

We attempted to use chromatin immunoprecipitation (ChIP) experiments with an epitope tagged allele of Rpd3 in mitotic cells to advance our model and were not able to attain reproducible Rpd3 chromatin association. Recent findings show that Rpd3 chromatin association is developmentally regulated and becomes much more robust and focused in haploid “Q” cells that have depleted their growth media and entered quiescence (McKnight *et al*. 2015). Perhaps relatedly, Rpd3 has prominent roles in meiotic development during which cells similarly transition into transcriptional dormancy in response to nutrient starvation (Xu *et al*. 2012; Yeheskely-Hayon *et al*. 2013; Lardenois *et al*. 2015). It seems plausible that the H3K4me0/H3K36me Rpd3S pathway we have identified using the sensitized Spt6-Spn1 TS allele background may have more relevant roles within these developmental contexts? It will be of interest to use more penetrating methods such as ChIP-Seq to advance our model in meiotic and/or Q cells.

By what molecular mechanisms might H3K4me (and H3K36me) impact Rpd3S function? Histone modifications are typically suggested to impact chromatin effector complex function through their purported roles in recruitment of these complexes to specific regions of chromatin where these modifications reside. While considerable evidence exists for this, in the case of Rpd3S, focused studies suggest that the histone binding activities of Eaf3 and Rco1 play little/no role in chromatin recruitment and, rather, impact Rpd3S activity on chromatin allosterically (Drouin *et al*. 2010). Indeed, more recent findings illuminate a similar scenario in the function of the acetyltransferase complex NuA4, whose enzymatic action on chromatin is controlled through subunits with defined histone binding specificities that have no apparent role in chromatin recruitment of NuA4 (Steunou *et al*. 2016). It is thus attractive to speculate that H3K4me3 opposition of Rco1 histone binding prevents Rpd3S from deacetylating nucleosomes through allosteric inhibition of Rpd3S activity (as opposed to through Rpd3S chromatin association). Whether H3K4/36me regulation of Rpd3S acts through chromatin recruitment or allosteric modulation of Rpd3 activity (or both), our model predicts Rpd3S deacetylation should be restricted near the 5’ ends of transcription units by H3K4me3, where this modification is typically found.

## Acknowledgements

This work was supported by CIHR grant MOP-89996 to M.D.M. We are grateful to Dr. Charlie Boone and Dr. Brenda Andrews for access to their temperature-sensitive mutant library and other reagents, and to Ziyan Chen for construction of Figure 3C. The *Rco1-TAP::HIS3MX6* and *Rco1*-Δ*PhD-TAP:HIS3MX6* strains were kindly provided by Dr. Jerry Workman.

---

**Supp. Figure 1.**
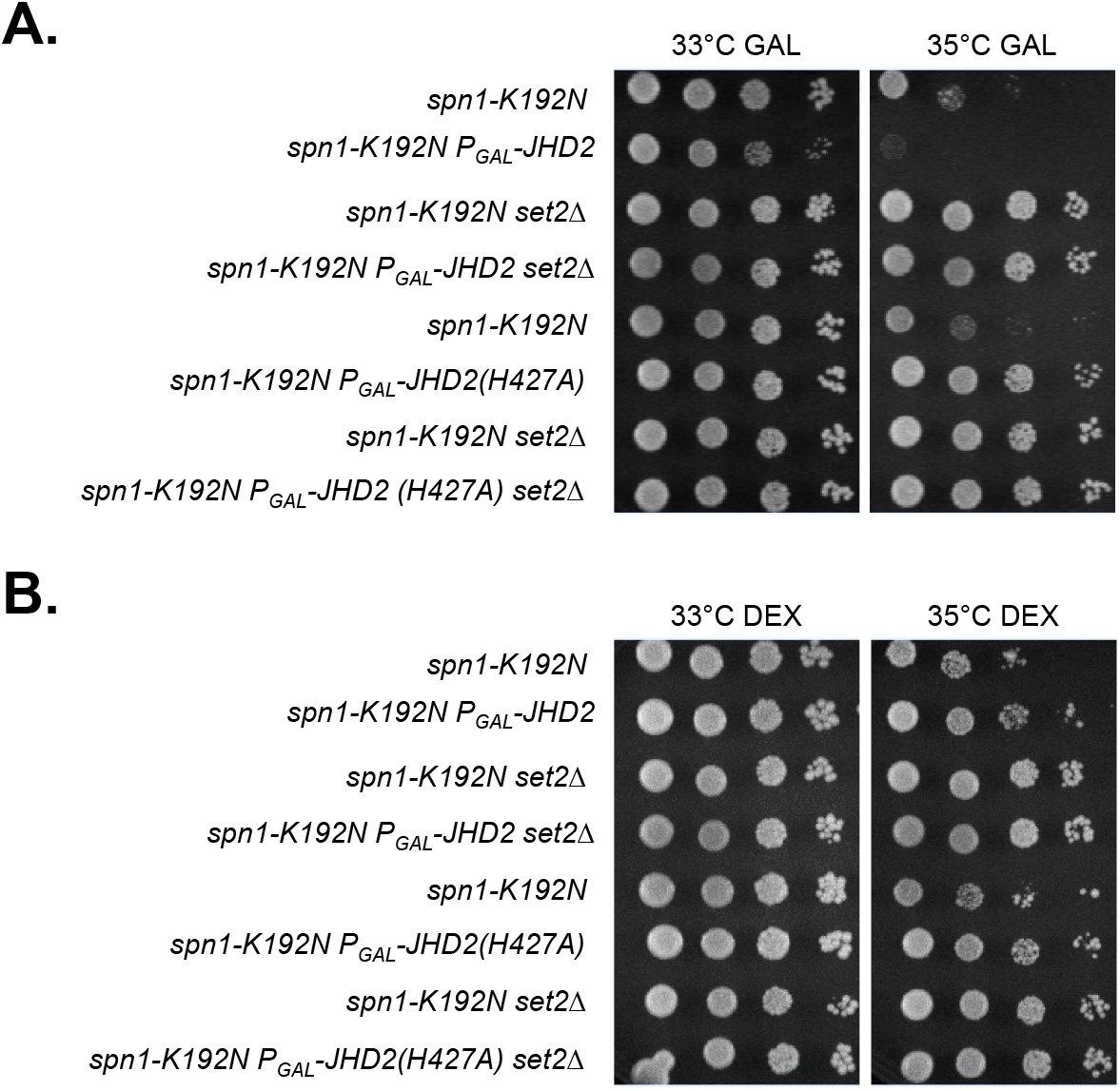
The enhancement of *spn1-K192N* by *JHD2* overexpression is reverted by a catalytically inactive histidine-427 to alanine mutation in Jhd2. Plate spot assays (as described in Figure 1) were used to compare the growth of the indicated strains on synthetic media with 2% dextrose (DEX) or 2% galactose (GAL). *P_GAL_-JHD2* is used to replace the endogenous *JHD2* locus so that cells grown in galactose overexpress *JHD2* and cells grown without galactose do not express JHD2. A H427A mutation was introduced into *P_gal_-JHD2* which disrupts its histone demethylase activity. (A) Genetic interactions of *set2*Δ and *P_GAL_-JHD2(H427A)* with *spn1-K192N*on GAL. (B) Genetic interactions of *set2*Δ and *P_GAL_-JHD2(H427A)* with *spn1-K192N*on DEX.

**Supp. Figure 2.**
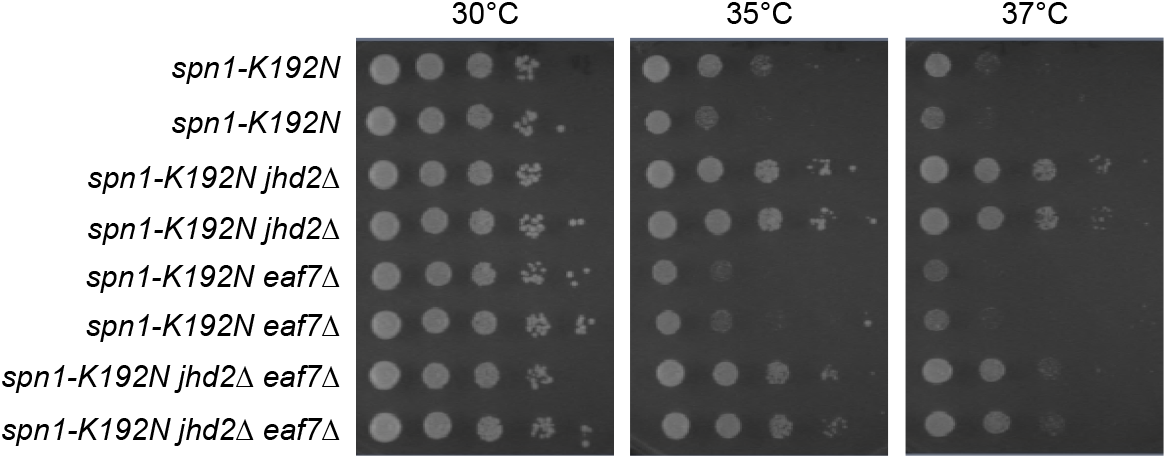
The suppression by *eaf3*Δ is not due to loss of the NuA4 histone acetyltransferase complex. Plate spot assays (as described in Figure 1) were used to compare the growth of the indicated strains. Genetic interactions of *eaf7*Δ and *jhd2*Δ with *spn1-K192N*.

**Supp. Figure 3.**
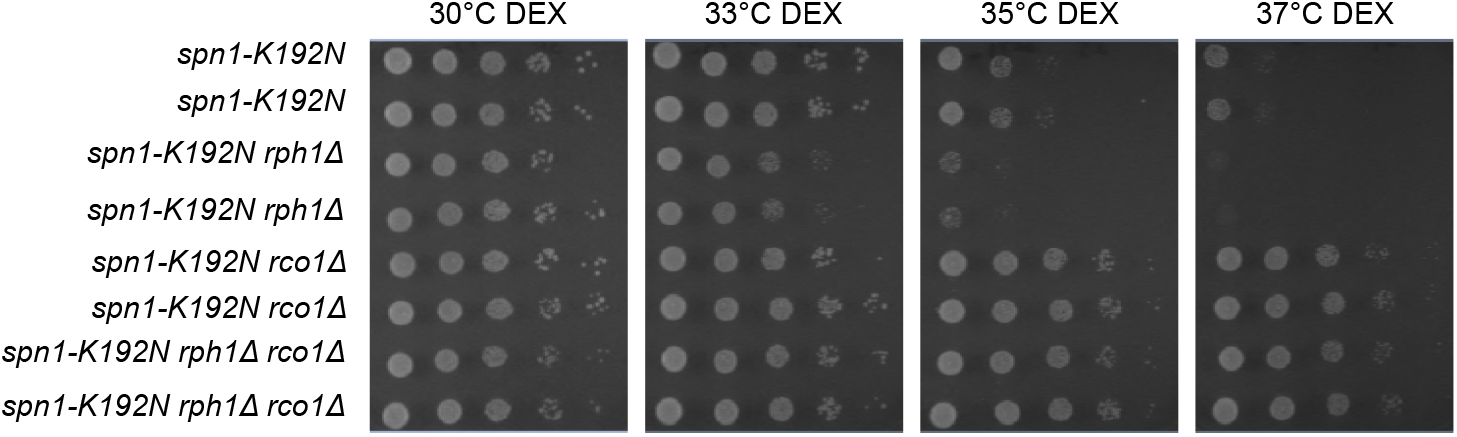
The enhancement of *spn1-K192N* temperature sensitivity by *rph1*Δ is reverted by *rco1*Δ. Plate spot assays (as described in Figure 1) were used to compare the growth of the indicated strains. Genetic interactions of *rcol*Δ and *rphl*Δ with *spn1-K192N*.

